# TERRA transcription destabilizes telomere integrity to initiate break-induced replication in human ALT cells

**DOI:** 10.1101/2021.02.18.431840

**Authors:** Bruno Silva, Rajika Arora, Claus M. Azzalin

## Abstract

Alternative Lengthening of Telomeres (ALT) is a Break-Induced Replication (BIR)-based mechanism elongating telomeres in a subset of human cancer cells. While the notion that spontaneous DNA damage at telomeres is required for ALT to occur, the molecular triggers of this physiological telomere instability are largely unknown. We previously proposed that the telomeric long noncoding RNA TERRA may represent one such trigger; however, given the lack of tools to suppress TERRA transcription in cells, our hypothesis remained speculative. We have now developed Transcription Activator-Like Effectors able to rapidly inhibit TERRA transcription from multiple chromosome ends in an ALT cell line. TERRA transcription inhibition decreases marks of DNA replication stress and DNA damage at telomeres and impairs ALT activity and telomere length maintenance. We conclude that TERRA transcription actively destabilizes telomere integrity in ALT cells, thereby initiating BIR and supporting telomere elongation. Our data point to TERRA transcription manipulation as a potentially useful target for therapy.

## INTRODUCTION

Transcription of telomeric DNA into the long noncoding RNA TERRA is an evolutionarily conserved feature of eukaryotic cells with linear chromosomes^1^. RNA polymerase II produces TERRA proceeding from subtelomeric regions towards chromosome ends and using the C-rich telomeric strand as a template. As a result, TERRA molecules comprise chromosome-specific subtelomeric sequences followed by a variable number of telomeric UUAGGG repeats^2-4^. TERRA is found either dispersed throughout the nucleoplasm or associated with telomeric chromatin, as well as other genomic loci that contain or not telomeric DNA repeats^4-7^. The molecular mechanisms mediating TERRA retention on chromosomes still need to be fully elucidated; however, the propensity of TERRA to form RNA:DNA hybrids with its template DNA strand (telomeric R-loops or telR-loops)^8-11^ and the physical interaction of human TERRA with the shelterin factors TRF1 and TRF2^12,13^ suggest that TERRA association with telomeric DNA-containing loci involves RNA/DNA and RNA/protein interactions.

The chromosomal origin of human TERRA is controversial. Using RT-PCR and Illumina sequencing, independent laboratories reported on the existence of TERRA molecules originating from a multitude of chromosome ends^3,14-19^. Consistently, we previously identified CpG dinucleotide-rich tandem repeats of 29 bp displaying promoter activity and located on approximately half of chromosome ends^2^. 29 bp repeats are positioned at variable distances from the first telomeric repeat and their transcriptional activity is repressed by CpG methylation^2^. Moreover, transcription factor binding sites exist on multiple subtelomeres and inactivation of some of them alter TERRA levels in cells^14,20-22^. However, work from the Blasco laboratory, based on reanalysis of TERRA Illumina sequencing and molecular and cell biological validation experiments, posed that human TERRA is mainly transcribed from one unique locus on the long arm of the chromosome 20 (20q) subtelomere^23,24^. The same group used CRIPSR/Cas9 to delete a 8.1 kb fragment from the 20q subtelomere comprising 4 putative promoters in U2OS osteosarcoma cells and isolated several clonal lines (20q-TERRA KO cells). Seemingly supporting the proposed origin of TERRA, 20q-TERRA KO cells displayed substantially diminished total TERRA levels when compared to parental cells^23,24^.

TERRA is involved in several telomere-associated processes including telomerase recruitment and regulation, telomeric DNA replication, telomeric heterochromatin establishment, response to DNA damage at telomeres and replicative senescence establishment^1^. Our laboratory and others have implicated TERRA also in telomere elongation in telomerase-negative cancer cells with an activated Alternative Lengthening of Telomeres (ALT) mechanism^8,9,24,25^. ALT is a specialized pathway repairing and thus re-elongating damaged telomeres through Break-Induced Replication (BIR) occurring in the G2 and M phases of the cell cycle and requiring the DNA polymerase delta accessory subunits POLD3 and POLD4^25-29^. Consistent with a function for TERRA in ALT, human ALT cells, including U2OS, are characterized by elevated telomeric transcription and TERRA levels, in part owed to hypomethylation of 29/37 repeats, and abundant telR-loops^2,8,15^. Moreover, the RNA:DNA endoribonuclease RNaseH1 and the ATPase/helicase FANCM dismantle telomeric R-loops and FANCM restricts total TERRA levels specifically in ALT cells. Because RNaseH1 and FANCM inactivation increases telomere instability and ALT activity, while their over-expression alleviates ALT^8,9,30,31^, we proposed that a physiological damage triggered by TERRA/telR-loops at ALT telomeres may provide the substrate for BIR-mediated telomere elongation^8,9,32,33^. However, due to a lack of tools to rapidly suppress TERRA transcription in cells, our hypothesis remained speculative. Further challenging our hypothesis, 20q-TERRA KO cells show increased telomeric localization of the DNA damage factors γH2AX and 53BP1 and telomeric fusions^24^, which has been interpreted as evidence for TERRA capping, rather than destabilizing, telomeres in ALT cells.

## RESULTS

### Development of Transcription Activator-Like Effectors binding to 29 bp repeats

To assess the short-term impact of TERRA transcription on telomere stability in ALT cells we engineered Transcription Activator-Like Effectors (TALEs)^34^ targeting a 20 bp sequence within the 29 bp repeat consensus^2^ (herein referred to as T-TALEs; Fig. 1a). Variable numbers of exact 20 bp sequences are found within the last 3 kb of 20 different subtelomeres (3p, 5p, 9p, 12p, 16p, 19p, 1q, 2q, 4q, 5q, 6q, 10q, 11q, 13q, 15q, 16q, 19q, 21q, 22q, and Xq/Yq). T-TALEs were C-terminally fused to a strong nuclear localization signal (NLS), four transcription repressor domains of the mSIN3 interaction domain (Enhanced Repressor Domain, SID4X) and a human influenza hemagglutinin (HA) epitope (Fig. 1a). T-TALEs not fused to SID4X were used as controls. Transgenes were cloned downstream of a doxycycline (dox) inducible promoter and stably integrated into U2OS cells expressing the tetracycline repressor protein. Several clonal cell lines were isolated in absence of dox and successively tested for dox-induced T-TALE expression by western blot and indirect immunofluorescence (IF) using anti-HA antibodies. Two independent cell lines for SID4X-fused T-TALEs (sid1 and sid4) and two for unfused T-TALEs (nls1 and nls3) were chosen for further experiments because: i) transgene expression was almost undetectable in absence of dox; and ii) dox treatments induced expression of ectopic proteins homogeneously distributed across the cell population, properly localized to the nucleus and at fairly similar levels in the four cell lines (Fig. 1a, S1a and b).

**Figure 1:**
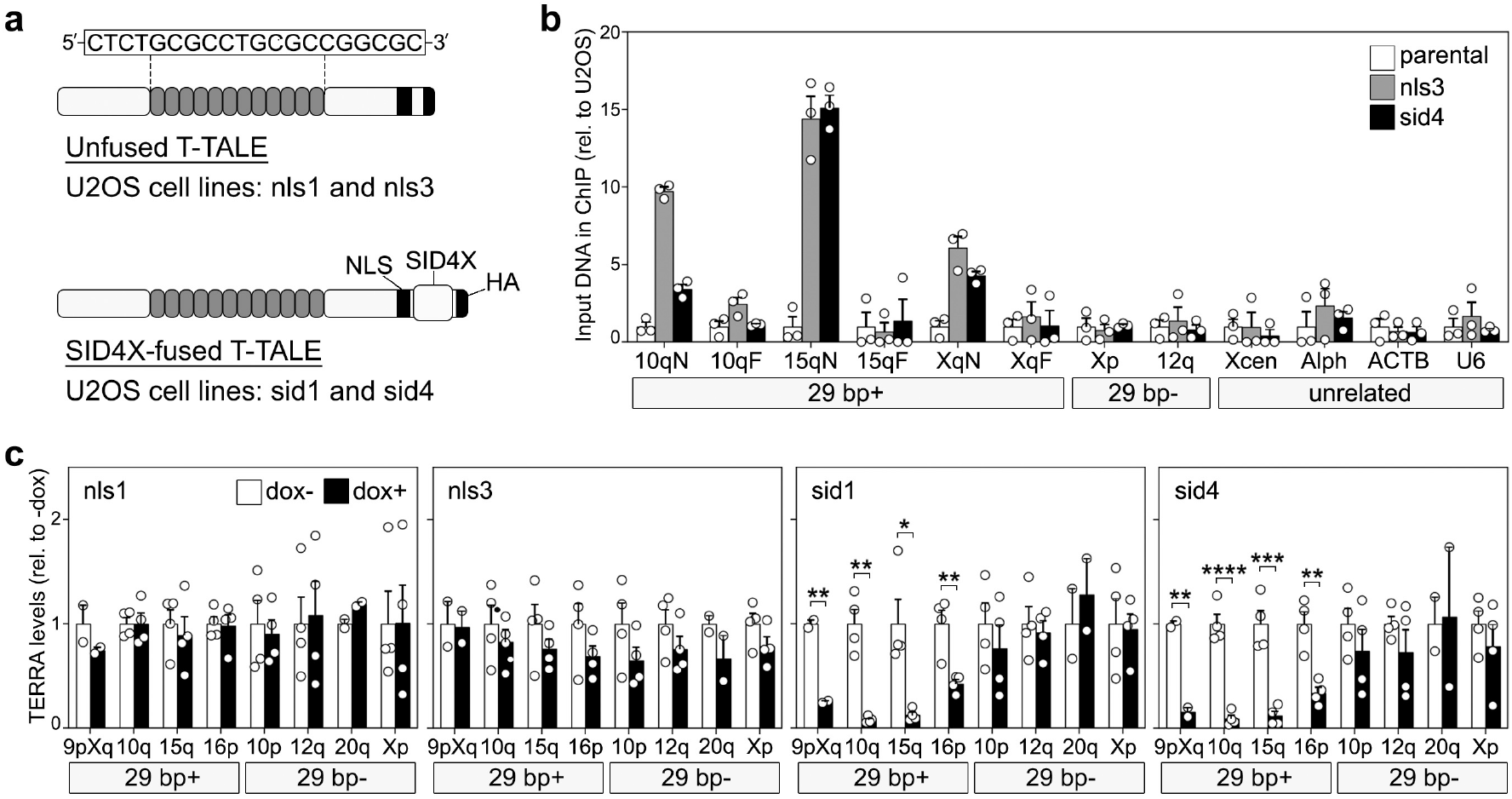
Development and validation of T-TALEs. (**a**) Schematic representation of T-TALEs. The RVD domain recognizing the indicated nucleotides within the 29 bp repeat consensus sequence is represented by grey rounded rectangles. NLS: nuclear localization signal; SID4X: four transcription repressor domains of the mSIN3 interaction domain; HA: human influenza hemagglutinin tag. (**b**) Quantification of anti-HA ChIPs in the indicated cell lines treated with dox for 24 hours. QPCRs were performed with oligonucleotides amplifying subtelomeric regions from chromosome ends containing or devoid of 29 bp repeats (29 bp+ and 29 bp-, respectively). For 29 bp+ subtelomeres, two oligonucleotide pairs placed at different distances from the 29 bp array were used and are indicated as N (near) and F (far). Control qPCR were performed with oligonucleotides amplifying sequences from a unique region of the X chromosome centromere (Xcen), alphoid DNA (Alph) and beta Actin (ACTB) and U6 gene loci. Values are graphed as input DNA found in the corresponding ChIP samples normalized to U2OS parental samples. Bars and error bars are means and SEMs from 3 independent experiments. Circles are single data points. (**c**) RT-qPCR quantifications of TERRA transcripts from 29 bp+ and 29 bp-chromosome ends in the indicated cell lines, treated with dox for 24 hours or left untreated. Values are graphed normalized to -dox. Bars and error bars are means and SEMs from 2 independent experiments for 9pXq and 20q and from 4 independent experiments for the remaining chromosome ends. Circles are single data points. *P* values were calculated with a two-tailed Student’s *t*-test. \**P* < 0.05, \*\**P* < 0.005, \*\*\**P* < 0.001, \*\*\*\**P* < 0.0001.

To confirm T-TALE binding specificity, we treated nls3, sid4 and parental U2OS cells with dox for 24 hours and performed chromatin immunoprecipitations (ChIPs) with anti-HA antibodies followed by qPCR using oligonucleotide amplifying subtelomeric sequences from several chromosome ends either containing or not 29 bp repeat sequences (29 bp+ and 29 bp-, respectively). DNA in very close proximity of 29 bp repeats on chromosomes 10q, 15q and XYq subtelomeres was enriched in nls3 and sid4 ChIP samples over parental U2OS samples. The enrichment diminished for sequences more distant from the 29 bp repeats on the same subtelomeres (Fig. 1b). On the contrary, no enrichment was observed for DNA from 29 bp-subtelomeres (XYp and 12q), the centromere of the X chromosome, centromeric alphoid repeats and the beta Actin or U6 gene loci (Fig. 1b). This confirms that T-TALEs specifically bind to 29 bp repeat.

### T-TALEs rapidly suppress TERRA transcription from 29 bp repeat-containing chromosome ends

To test the functionality of T-TALEs, we treated cells with dox for 24 hours and performed RT-qPCR to measure TERRA levels from 29 bp+ (9Xq, 10q, 15q and 16p) and 29 bp-(10p, 12q, 20q and XYp) subtelomeres. In sid1 and sid4 cells, TERRA from 29 bp+ subtelomeres was greatly reduced while TERRA from 29 bp-was not affected. In nls1 and 3 cells, no significant change in TERRA levels was observed at any chromosome ends (Fig. 1c). Fluorescence-activated cell sorting (FACS) of propidium iodide (PI)-stained cells did not reveal cell cycle profile alterations in dox-treated nls and sid cells as compared to untreated controls (Fig. S2a and b); this indicates that the TERRA decrease observed in sid1 and sid4 cells is not an indirect consequence of a disturbed cell cycle progression. Further supporting this notion, in ALT cells, TERRA levels do not diminish when the cell cycle progresses from S to G2 phases, as typical in telomerase positive cells^4,35^. Hence, our TALE-based system can efficiently and specifically inhibit TERRA transcription from 29 bp promoter repeats. Notably, because TERRA transcription from 29 bp-chromosome ends is not affected in dox-treated sid cells, an immediate cross-talk between the transcriptional state of independent telomeres appears not to exist in U2OS cells. However, a larger number of ends would need to be tested to corroborate this conclusion.

### TERRA transcription suppression alleviates telomere instability

To probe the effects of TERRA transcription inhibition on telomere stability, we performed indirect immunofluorescence (IF) using antibodies against the single-stranded DNA binding protein RPA32, RPA32 phosphorylated at serine 33 (pSer33) or γH2AX combined with either telomeric DNA fluorescence in situ hybridization (FISH) or IF against the shelterin component TRF2. RPA32 and pSer33 were used as markers of DNA replication stress, while γH2AX as a broad DNA damage marker. Dox treatments diminished the telomeric localization of both RPA32 variants in sid1 and sid4 cells (Fig. 2a and b, Fig. S3) already at 24 hours after drug delivery. A slightly sharper decrease was observed for total RPA32 than for pSer33, suggesting that a fraction of the protein binds to telomeres independently of serine 33 phosphorylation or that the protein undergoes dephosphorylation while still telomere-bound. Similarly, dox treatments diminished the frequencies of γH2AX co-localization with telomeres in sid1 and sid4 cells (Fig. 2a and b, Fig. S3). Dox treatments did not alter co-localization frequencies for any of the tested markers in nls1 and nls3 cells (Fig. 2a and b, Fig. S3). Moreover, dox did not affect the total cellular levels of RPA32, pSer33, γH2AX and TRF2 nor did it impair RPA32 and H2AX phosphorylation when cells were simultaneously treated with the damaging agent camptothecin (Fig. S1b). Hence, all changes observed in dox-treated sid1 and sid4 cells do not derive from altered protein cellular levels or from a compromised DNA damage response. Alterations in cell cycle distribution also cannot account for the observed changes in RPA32, pSer33 and γH2AX at telomeres (Fig. S2a and b).

**Figure 2:**
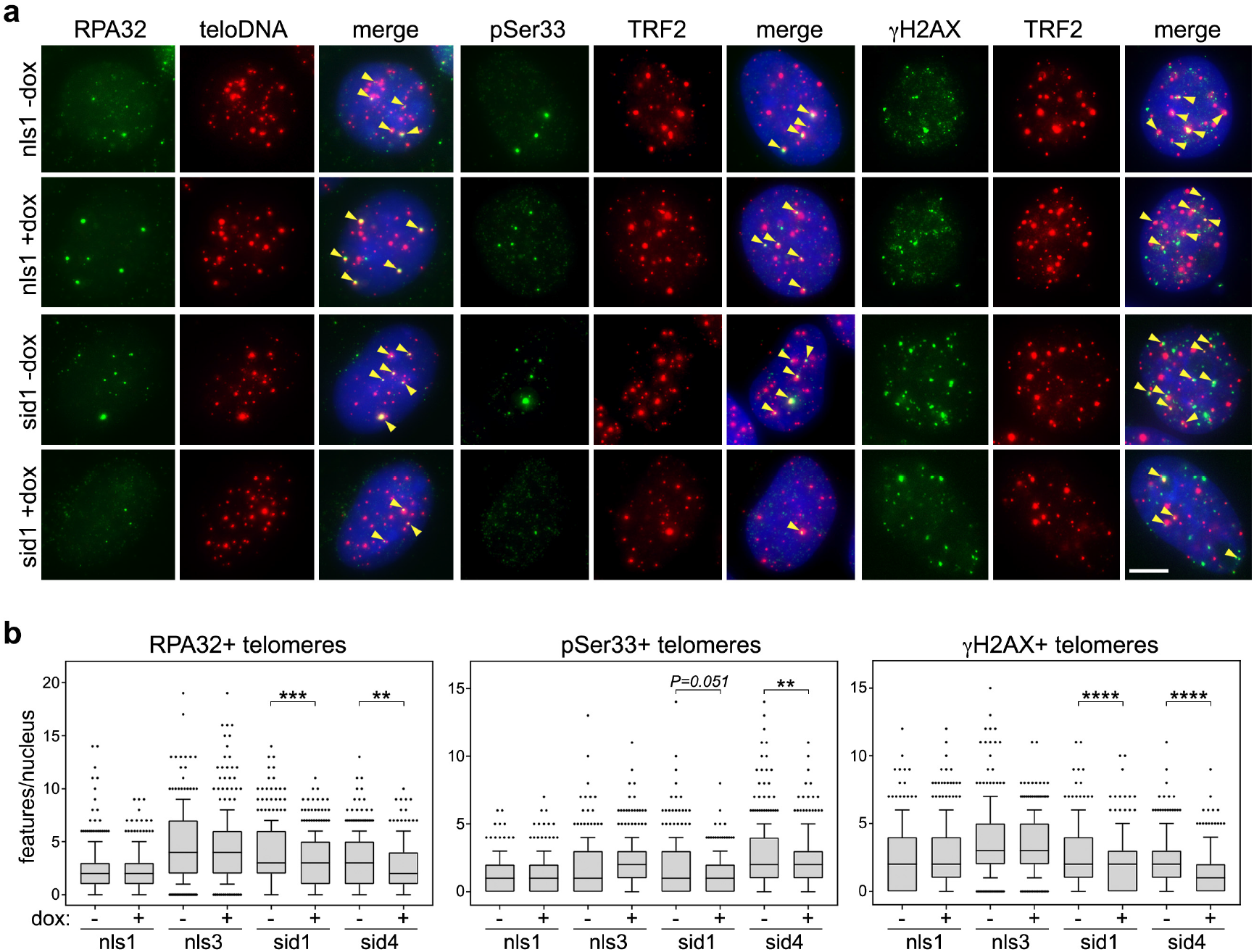
TERRA transcription inhibition alleviates telomere instability. (**a**) Examples of RPA32, pSer33 or γH2AX IF (green) combined with telomeric DNA FISH (teloDNA) or TRF2 IF (red). DAPI stained DNA is in blue. The indicated cell lines were treated with dox for 24 hours for RPA32 and pSer33 or 72 hours for γH2AX. Arrowheads in the merge panels point to co-localization events. Scale bar: 5 μm. (**b**) Box plots of the 10-90 percentile of co-localization events per nucleus in experiments as in **a**. Central lines are medians. A total of at least 300 nuclei from three independent experiments were analyzed for each sample. *P* values were calculated with a Mann-Whitney *U* test. \*\**P* < 0.005, \*\*\**P* < 0.001, \*\*\*\**P* < 0.0001.

### TERRA transcription suppression inhibits ALT activity and telomere maintenance

According to our model, diminished telomere instability should weaken ALT activity. To test this, we first quantified ALT-associated PML bodies (APBs) by combining IF with an anti-PML antibody and telomeric DNA FISH. APBs diminished in sid1 and sid4 but not in nls1 and nls3 cells already 24 hours after adding dox (Fig. 3a and b, Fig. S3). We then synchronized cells at the G1/S transition and let them progress from S-phase to G2 in presence of dox and the Cdk1 inhibitor RO-3306. Cells were pulsed with EdU during the last 2.5 hours of treatment and subjected to EdU detection combined with telomeric DNA FISH. Dox did not affect the frequencies of EdU co-localization with telomeric DNA in nls1 and nls3 cells, while it substantially diminished them in sid1 and sid4 cells (Fig. 3a and b, Fig. S3). This suggests that dox treatments reduced telomeric BIR is in G2 synchronized sid cells. Consistently, as shown by double IF experiments, dox diminished the frequencies of POLD3 co-localization with the shelterin component RAP1 in sid1 and sid4, but not in nl1 and nls3 G2 cells (Fig. 3a and b, Fig. S3). Changes in APBs and POLD3 telomeric localization occurred in absence of changes in PML, POLD3 and RAP1 total protein levels (Fig. S1b). Moreover, dox treatments did not affect the efficiency of our synchronization protocol (Fig. S2a and b). Thus, the decline in ALT features observed in sid1 and sid4 cells cannot be ascribed to altered protein levels or differences in the fraction of cells in G2 phase.

**Figure 3:**
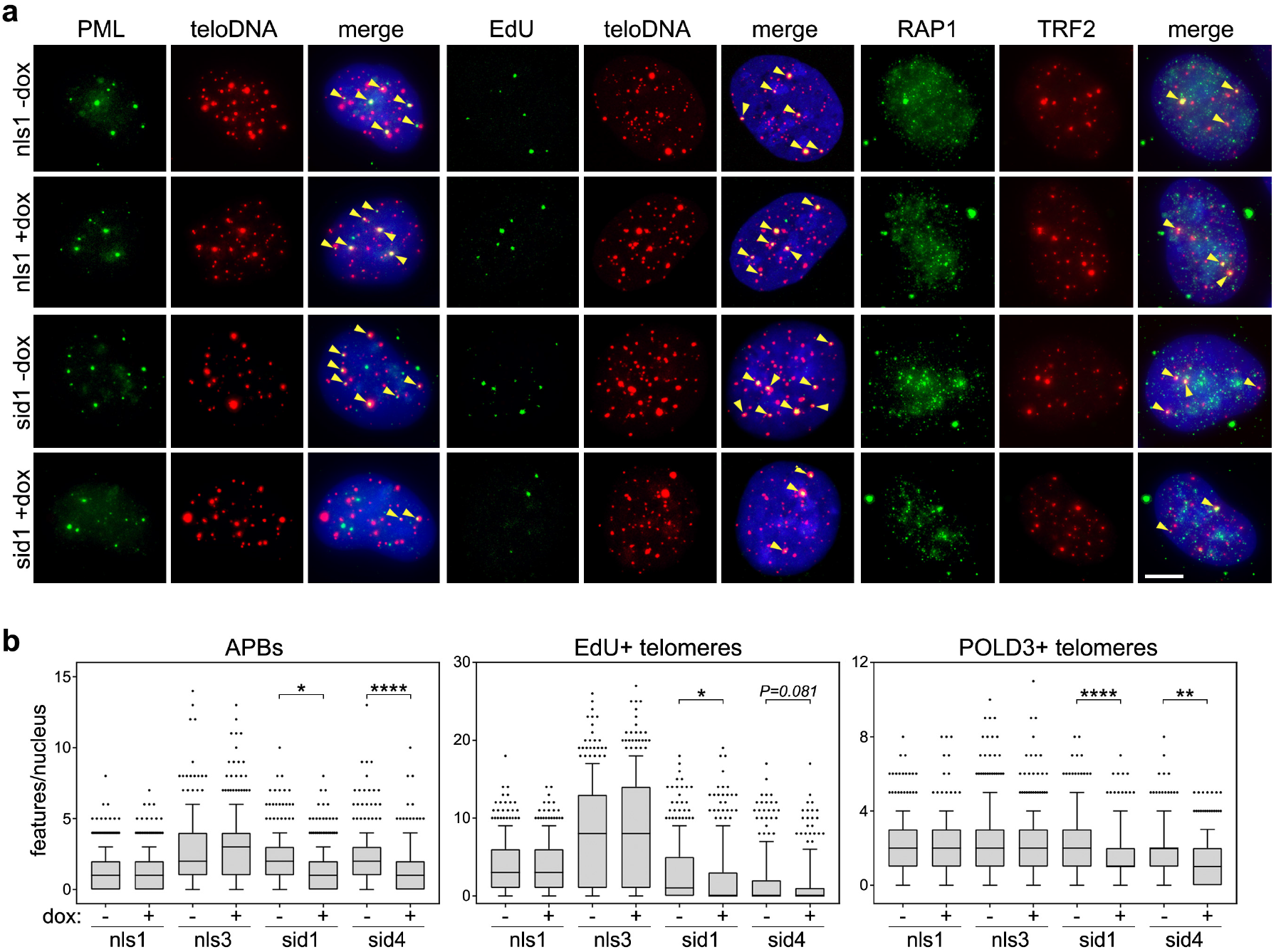
TERRA transcription inhibition alleviates ALT activity. (**b**) Left and right panels: examples of PML or POLD3 IF (green) combined with telomeric DNA FISH (teloDNA) or RAP1 IF (red). Middle panels: examples of EdU detection (green) combined with telomeric DNA FISH (red). DAPI stained DNA is in blue. The indicated cell lines were treated with dox for 24 hours for PML or for 24.5 hours for POLD3 and EdU (G2/M synchronized cells, see methods for details). Arrowheads in the merge panels point to co-localization events. Scale bar: 5 μm. (**b**) Box plots of the 10-90 percentile of co-localization events per nucleus in experiments as in **a**. Central lines are medians. A total of at least 300 nuclei from three independent experiments were analyzed for each sample. *P* values were calculated with a Mann-Whitney *U* test. \**P* < 0.05,\*\**P* < 0.005, \*\*\*\**P* < 0.0001.

As additional markers for ALT activity, we also quantified C-circles, which are circular telomeric DNA molecules with exposed single stranded C-rich tracts, and telomeric sister chromatid exchanges (TSCEs)^26^. C-circle assays with total genomic DNA did not disclose consistent changes in sid1, nls1 and sid4 cells treated with dox for up to 72 hours. In nls3 cells, C-circle levels diminished after 24 hours of treatment and started to recover at later timepoints (Fig. S4a and b). Chromosome orientation FISH (CO-FISH) on nls3 and sid4 metaphase chromosomes did not detect significant changes in TSCE frequencies associated with dox treatments in either cell line (Fig. 4a-c). However, we noticed unequal distribution of leading and lagging strand signals at several chromosome ends. Thus, we also quantified the occurrence of sister telomeres with 2 leading and one lagging strand signals (double leading or DLead) and with 2 lagging and one leading strand signals (double lagging or DLagg). DLagg telomeres were not affected by dox treatments in both cell lines, while DLead telomeres were more than halved in sid4 but not nls3 dox-treated cells as compared to untreated controls (Fig. 4a-c).

**Figure 4:**
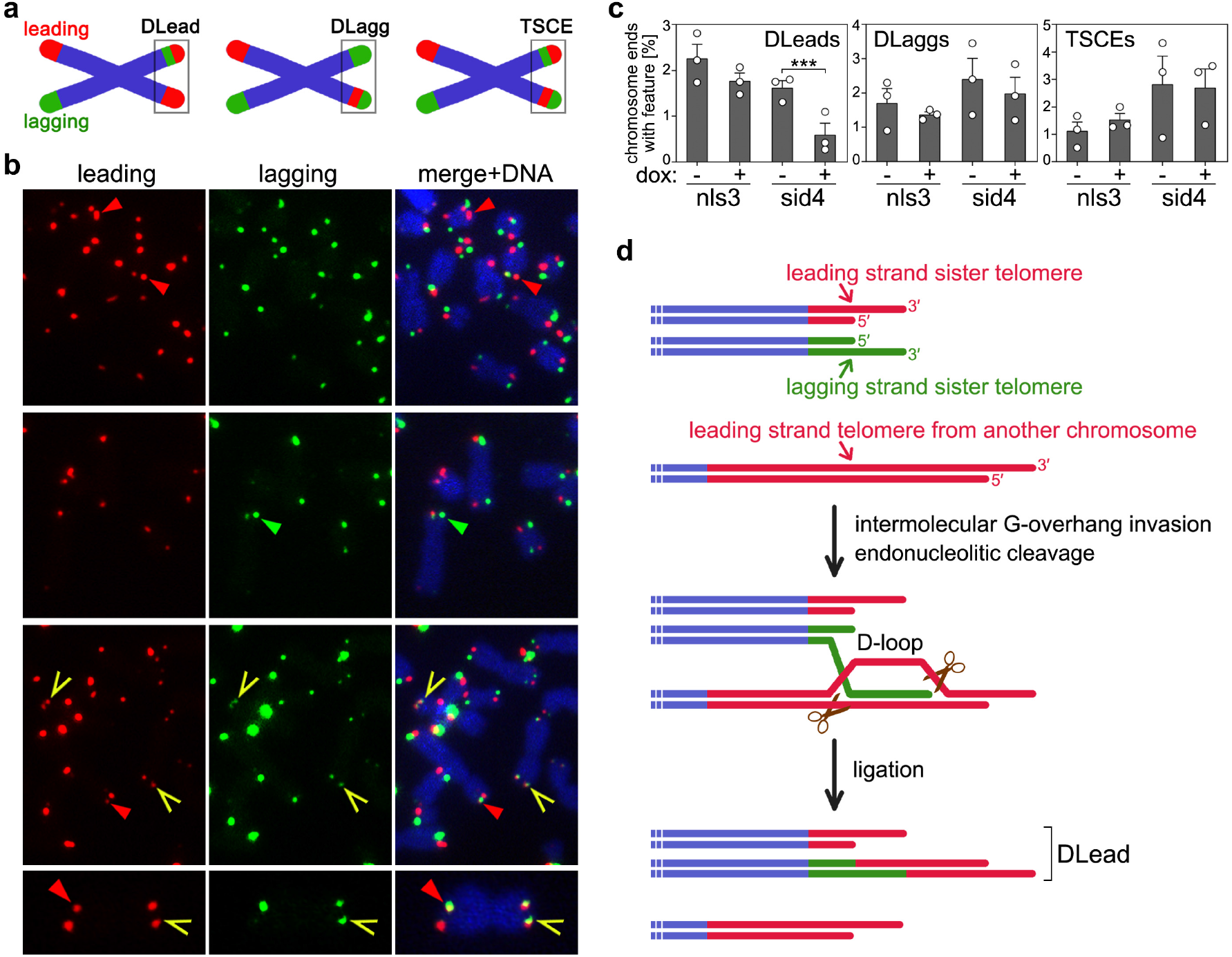
TERRA transcription inhibition diminishes the frequencies of DLead chromosome ends. (**a**) Schematic representation of telomeric features scored in CO-FISH experiments. DLead: sister telomeres with 2 leading and one lagging strand signals; DLagg: sister telomeres with 2 lagging and one leading strand signals; TSCEs: telomeric sister chromatid exchanges. (**b**) Examples of CO-FISH on metaphases from sid4 cells treated with dox for 72 hours. Leading and lagging strand telomeres are in red and green, respectively; DAPI stained DNA is in blue. Red arrowheads point to DLeads, green arrowheads to DLaggs and yellow arrows to TSCEs. A chromosome with one DLead and one TSCE at its two opposite ends is shown at higher magnification at the bottom. (**c**) Quantifications of telomeric features in CO-FISH experiments as in **b**. A total of at least 2538 chromosomes from 3 independent experiments were analyzed for each condition. Bars and error bars are means and SEMs. Circles are single data points. *P* values were calculated with a two-tailed Student’s *t*-test. \*\*\**P* < 0.001. (**d**) Speculative model for DLead generation. Scissors represent structure specific endonucleases. See Discussion for details.

Alleviation of ALT activity should translate into impaired telomere elongation and progressive loss of telomeric DNA. We treated cells with dox over a prolonged time course and analyzed telomeres by telomeric DNA FISH on metaphase chromosomes. TERRA transcription suppression in sid 1 and sid4 cells was maintained throughout the entire time course (data not shown). A progressive, statistically significant accumulation of telomere free ends (TFEs) was observed in sid1 and sid4 cells treated with dox for up to 15 and 9 days, respectively (Fig. 5a and b). Conversely, no significant change in TFEs was observed in nls3 cells during the tested time course (Fig. 5a and b).

**Figure 5:**
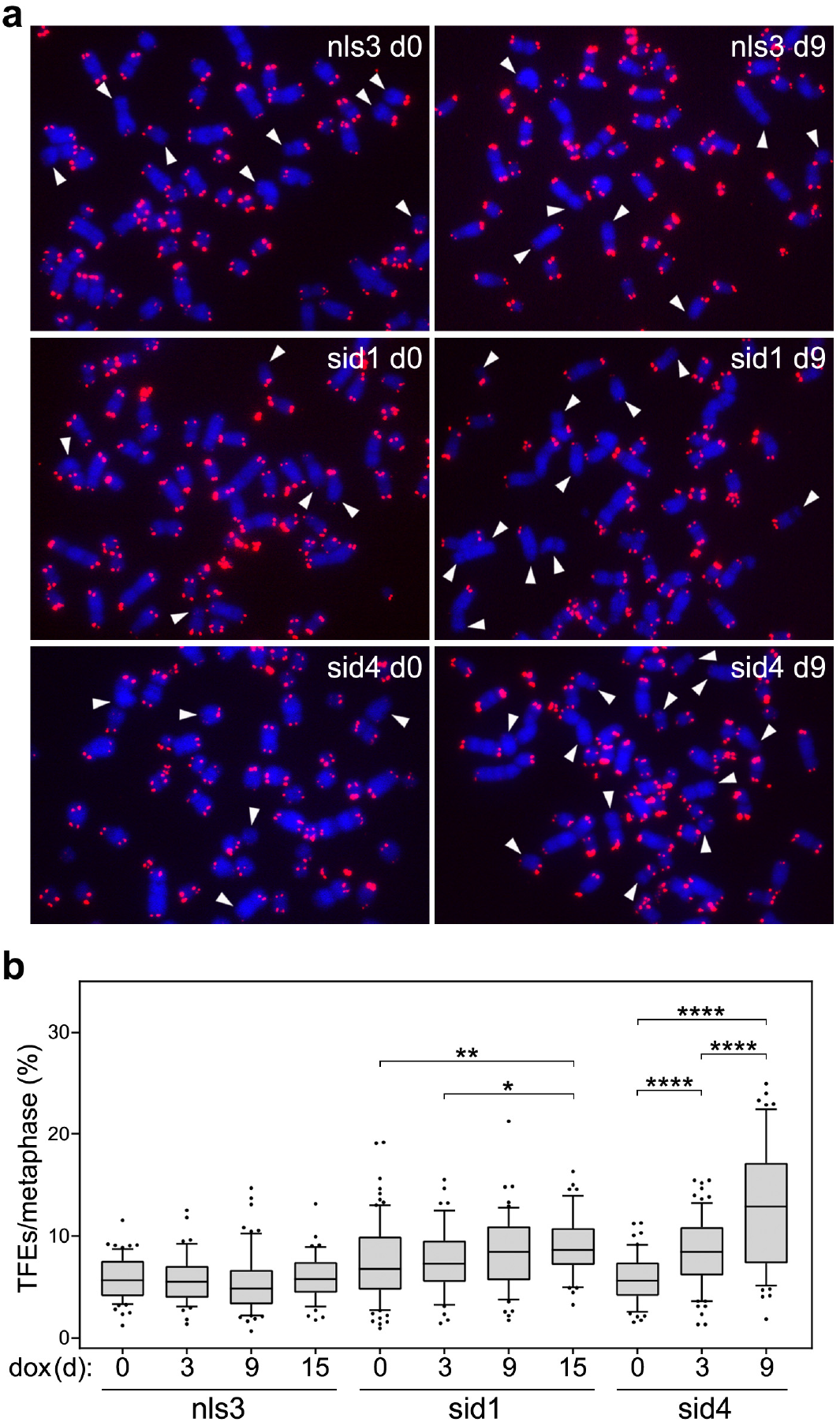
TERRA transcription inhibition leads to accumulation of TFEs. (**a**) Examples of telomeric DNA FISH on metaphases from the indicated cell lines treated with dox for 0 or 9 days (d). Telomeric repeat DNA is in red, DAPI-stained chromosomal DNA in blue. White arrowheads point to chromosome arms with TFEs. (**b**) Box plots of the 10-90 percentile of TFEs per metaphase in experiments as in **a**. Central lines are medians. Cells were harvested at the indicated days of dox treatment. A total of at least 2761 chromosomes from 2 or 3 independent experiments were analyzed for each condition. *P* values were calculated with a Mann-Whitney *U* test. \**P* < 0.05, \*\**P* < 0.005, \*\*\*\**P* < 0.0001.

## DISCUSSION

We have developed an efficient system to suppress TERRA transcription from several chromosome ends in an ALT cell line. Importantly, rapid suppression of TERRA transcription across a cell population provides the critical advantage to study the immediate consequences on telomere homeostasis, avoiding secondary effects associated with clonal selection and expansion after TERRA inhibition. Our data clearly establish that TERRA transcription inhibition impairs accumulation of DNA instability markers (RPA32 and γH2AX) at telomeres, weakens ALT features (APBs and PLD3-dependent synthesis of telomeric DNA in G2 cells) and causes TFE generation. We propose that, in ALT cells, TERRA transcription is a major trigger of replication stress-associated telomere instability and, in turn, BIR-mediated telomere elongation. The only partial decrease in telomere instability and ALT activity observed in dox-treated sid1 and sid4 cells is most likely explained by the fact that T-TALEs only target a subset of telomeres and/or the existence of additional telomere instability triggers, for example G-quadruplex structures^25^. Based on previous studies on RNAseH1 and FANCM^8,12^ and the ability of R-loops to induce DNA instability^36^, it seems likely that TERRA transcription causes replication stress by stalling the replication fork through telR-loop formation. It is also possible that telomere instability derives from the collision between TERRA transcription and telomeric replication forks. Importantly, our data argue against the notion that TERRA caps ALT telomeres, as it was previously proposed based on the massive accumulation of telomeric DNA damage in 20q-TERRA KO cells^24^. On the contrary, TERRA transcription actively destabilizes the integrity of ALT telomeres to support their elongation and cell immortality. In our view, the telomeric DNA damage detected in U2OS 20q-TERRA KO cells originates from the very short telomeres present in those cells and does not underscore direct TERRA-associated capping functions^24^.

It is interesting that not all tested ALT features, including C-circles, are affected when TERRA transcription is inhibited. Two alternative BIR pathways have been shown to co-exist in ALT cells, one RAD52-dependent and associated with C-circle production, and the other RAD52-independent and not leading to C-circle production^29^. It is possible that TERRA only supports RAD52-independent BIR, although this hypothesis is not consistent with the observed C-circle accumulation in RNaseH1-and FANCM-depleted ALT cells^8,9,30,31^. We thus consider that mild changes in C-circles upon partial TERRA transcription inhibition may fall below the detection limit of our assays. TSCEs are also not affected when TERRA transcription is inhibited. This observation was surprising because 20q-TERRA KO cells are characterized by diminished TSCE frequencies^24^. However, a substantial fraction of the overall short telomeres in 20q-TERRA KO cells might have escaped detection in CO-FISH experiments, thus skewing the results and their interpretation. Regardless, our analysis in T-TALE cells revealed that TERRA transcription promotes the formation of rearranged chromosome ends with leading strand replication DNA present at both sisters (DLeads); hence, the mechanisms leading to TSCEs and DLeads are different. Because TERRA transcription promotes telomeric BIR, we interpret DLead structures as the outcome of premature termination of post-replicative BIR events initiating with a lagging strand telomere invading a leading strand one from another chromosome (Fig. 4d). As soon as a D-loop is formed, it could be resolved by structure-specific endonucleases, for example the SMX complex, which has been previously implicated in ALT^37-41^. Endonucleolytic cleavage followed by end-joining would translocate the distal part of the leading strand telomere onto the lagging strand one, thus generating a DLead structure (Fig. 4d).

Events where lagging strand telomeres translocate onto leading strand ones, thereby generating DLagg ends, also seem to occur. However, because TERRA transcription inhibition does not affect DLagg frequencies, different molecular triggers appear to act at leading and lagging strand telomeres to initiate BIR. The specificity of TERRA transcription on DLead frequencies can be explained in two alternative, yet not mutually exclusive ways. One possibility is that TERRA transcription increases the propensity of lagging strand telomeres to invade other chromosome ends, for example by inducing replication fork stalling and DSBs through telR-loop formation^36^. Because telR-loops, at least in budding yeast, are more abundant at short telomeres^42^, this might constitute a regulatory mechanism directing ALT towards the shortest telomeres in the cell. Alternatively, TERRA transcription could prime leading strand telomeres to act as templates for BIR by altering their structure; this also could depend on the formation of telR-loops, as they might shape the double helix into an ideal entry platform for an annealing reaction involving a switch between TERRA and the 3’ end of the acceptor telomere. This second hypothesis is supported by the observation that, in ALT cells, aberrant telR-loop accumulation due to RNAseH1 depletion causes rapid loss of leading strand telomeres^8^.

Our findings also unmistakably settle that multiple chromosome ends, and not only the one of 20q, are actively transcribed and further validate that the previously identified 29 pb repeats are functional and physiologically relevant TERRA promoters^2^. We believe that the diminished levels of cellular UUAGGG repeats in 20q-TERRA KO cells^23,24^ derive from the short telomeres in those cells and/or clonal variability, and thus do not imply that 20q is the main TERRA locus in ALT cells. Moreover, although our T-TALEs do not affect 20q-TERRA transcription, which is expected because the 20q subtelomere does not contain 29 bp repeats, impaired telomere maintenance is observed both in our system and in 20q-TERRA KO cells; this suggests that many if not all telomeres in a cell have to stay transcriptionally active to assure proper ALT-mediated elongation.

Lastly, we anticipate that TERRA transcription might become a useful target for therapy. The physiological instability of ALT telomeres has to be kept within a precise range in order to trigger sufficient BIR yet without causing cell death^32^, implying that TERRA transcription must also be controlled. Decreasing TERRA transcription, for example through chemical inhibition of TERRA promoter activity, is expected to alleviate replication stress and lead to inefficient elongation and, in the long term, cell proliferation arrest. On the other hand, increasing TERRA transcription would generate excessive telomere instability, which, if above cellular tolerance, could cause sudden cell death. Modulating TERRA transcription holds the potential to spearhead unprecedented therapeutic protocols in the fight against ALT cancers.

## Acknowledgements

ACKNOWLEDGMENTS

We thank Harry Wischnewski for help with RT-qPCRs and C-circle assays, and the Bioimaging and Flow Cytometry facilities of iMM for services. This work was supported by the European Molecular Biology Organization (IG3576) and the Fundação para a Ciência e a Tecnologia (IF/01269/2015; PTDC/MED-ONC/28282/2017; PTDC/BIA-MOL/29352/2017).

## AUTHOR CONTRIBUTIONS

R.A. and C.M.A. conceived the original project. B.S., R.A. and C.M.A designed and performed the experiments, analyzed the data and wrote the manuscript.

## DATA AVAILABILITY

The authors declare that the data supporting the findings of this study are available within the paper and its supplementary information files. Plasmid sequences are available upon request.

## MATERIALS AND METHODS

### Plasmid construction

A repeat-variable di-residue (RVD) domain specifically targeting the 5’-CTCTGCGCCTGCGCCGGCGC-3’ sequence within the 29 bp repeat consensus sequence was designed using TAL Effector Targeter^43^. Variable numbers of the target 20 bp sequence are identified within the most distal 3 kb of 20 subtelomeres according to a complete clone-based assembly of human subtelomeric regions^44^. A 3560 bp DNA fragment corresponding to a complete TALE module comprising the designed RVD followed by an SV40 nuclear localization signal and a human influenza hemagglutinin (HA) tag was synthesized at GenScript. The fragment was cloned into KpnI/ApaI digested a pcDNA5-FRT-TO plasmid (ThermoFisher Scientific) downstream of a doxycycline-inducible CMV promoter (unfused T-TALE). The obtained plasmid was digested with ClaI and EcoRV and ligated to a 429 bp fragment comprising the Enhanced Repressor Domain and synthesized at GenScript (SID4X T-TALE). Plasmid sequences are available upon request.

### Cell culture procedures

T-TALE expressing U2OS cells were generated by FRT-mediated integration of unfused T-TALE and SID4X T-TALE plasmids into T-REx™-U2OS cells expressing the TetR protein (ThermoFisher Scientific). Clonal selection was performed by plating cells at low dilution in high glucose DMEM, GlutaMAX (Thermo Fisher Scientific) supplemented with 10% tetracycline-free fetal bovine serum (Pan BioTech), 100 U/ml penicillin-streptomycin (Thermo Fisher Scientific) and 200?μg/ml hygromycin B (VWR). Individual clones were manually picked and expanded in the same medium. For T-TALE induction, 50 ng/ml doxycycline (dox; Sigma-Aldrich) was added to the culture medium devoid of hygromycin B for 24-72 hours; for longer induction times, dox was refreshed every 72 hours. Mycoplasma contaminations were tested using the LookOut Mycoplasma PCR Detection Kit (Sigma-Aldrich) according to the manufacturer’s instructions. When indicated, cells were treated with 1 μM camptothecin (Sigma-Aldrich) for 6 h.

### Fluorescence-activated cell sorting (FACS)

Cells were trypsinized and pelleted by centrifugation at 500 g at 4°C for 5 min. Cell pellets were fixed in 70% ethanol at -20°C for 30 min and treated with 25 μg/ml RNaseA (Sigma-Aldrich) in 1x PBS at 37°C for 20 min. Cells were then centrifuged as above and pellets washed in 1x PBS and stained with 20 μg/ml propidium iodide (Sigma-Aldrich) in 1x PBS at 4°C for 10 min. Flow cytometry was performed on a BD Accuri C6 (BD Biosciences). Data were analyzed using FlowJo software.

### Western blotting

Cells were trypsinized and pelleted by centrifugation at 500 g at 4°C for 5 min. Pellets were resuspended in 2x lysis buffer (4% SDS, 20% Glycerol, 120 mM Tris-HCl pH 6.8), boiled at 95°C for 5 min and centrifuged at 1600 g at 4°C for 10 min. Supernatants were recovered and protein concentrations determined by Lowry assay using bovine serum albumin (Sigma-Aldrich) as a standard. 30 μg of proteins were mixed with 0.004% Bromophenol blue and 1% β-Mercaptoethanol (Sigma-Aldrich), incubated at 95°C for 5 min, separated in polyacrylamide gels, and transferred to nitrocellulose membranes (Maine Manufacturing, LLC) using a Trans-Blot SD Semi-Dry Transfer Cell apparatus (Bio-Rad). The following primary antibodies were used: a rabbit monoclonal anti-HA (Cell Signaling, 3724; 1:1000 dilution), a rabbit polyclonal anti-RAP1 (Bethyl, A300-306A; 1:2000), a mouse monoclonal anti-PCNA (Santa Cruz Biotechnology, sc-56; 1:10000), a rabbit polyclonal anti-TRF2 (Novus Biologicals, NB110-57130; 1:2000), a mouse monoclonal anti-POLD3 (Novus Biologicals, H00010714-M01; 1:500), a mouse monoclonal anti-ACTB (Santa Cruz Biotechnology, sc-47778; 1:2000), a mouse monoclonal anti-PML (Santa Cruz Biotechnology, sc-966; 1:500), a rabbit polyclonal anti-pSer33 (Bethyl, A300-246A; 1:2000), a rabbit polyclonal anti-RPA32 (Bethyl, A300-244A; 1:1000), a rabbit polyclonal anti-LMB1 (GeneTex, GTX103292S; 1:5000), a rabbit polyclonal anti-H3 (Santa Cruz Biotechnology, sc-10809; 1:4000), a mouse monoclonal anti-γH2AX (Millipore, 05-636, 1:2000), a rabbit polyclonal anti-H3 (Santa Cruz Biotechnology, sc-10809; 1:4000). Secondary antibodies were HRP-conjugated goat anti-mouse and anti-rabbit IgGs (Bethyl Laboratories, A90-116P and A120-101P; 1:3000). Signals were acquired using an Amersham 680 blot and gel Imager.

### DNA fluorescence *in situ* hybridization (FISH) and chromosome orientation FISH (CO-FISH)

Metaphase spreads were prepared by incubating cells with 200?ng/ml Colchicine (Sigma-Aldrich) for 5?h. Mitotic cells were harvested by shake-off and incubated in 0.075?M KCl at 37?°C for 10 min. Chromosomes were fixed in ice-cold methanol/acetic acid (3:1) and spread on glass slides. Slides were treated with 20 μg/ml RNase A (Sigma-Aldrich), in 1x PBS at 37?°C for 1?h, fixed in 4% formaldehyde (Sigma-Aldrich) in 1x PBS for 2 min, and treated with 70 μg/ml pepsin (Sigma-Aldrich) in 2 mM glycine, pH 2 (Sigma-Aldrich) at 37°C for 5 min. Slides were fixed again with 4% formaldehyde in 1x PBS for 2 min, incubated subsequently in 70%, 90% and 100% ethanol for 5 min each, and air-dried. A C-rich telomeric PNA probe (5’-Cy3-OO-CCCTAACCCTAACCCTAA-3’; Panagene) diluted in hybridization solution (10 mM Tris-HCl pH 7.2, 70% formamide, 0.5% blocking solution (Roche)) was applied onto the slides followed by incubation at 80°C for 5 min and at room temperature for 2 h. Slides were washed twice in 10 mM Tris-HCl pH 7.2, 70% formamide, 0.1% BSA and three times in 0.1 M Tris-HCl pH 7.2, 0.15 M NaCl, 0.08% Tween-20 at room temperature for 10 min each. For CO-FISH, cells were incubated with BrdU:BrdC (3:1, final concentration 10 μM; Sigma-Aldrich) for 16 h prior to metaphase preparation as above. Chromosomes were spread on glass slides, treated with RNaseA as above and incubated with 10 μg/ml Hoechst 33258 (Invitrogen) in 2x SSC for 15 min at room temperature. Slides were exposed to 365-nm ultraviolet light using a Stratagene Stratalinker 1800 UV irradiator set to 5400 Joules, and incubated with 3U/μl Exonuclease III (New England Biolabs) at 37°C for 30 min. Subsequent hybridizations were performed in 30% formamide, 2x SSC for 3h at room temperature using first a C-rich telomeric LNA probe (5’-6-FAM-CCCTAACCCTAACCCTAA-3’; Exiqon) and then a G-rich telomeric LNA probe (5’-TYE563-TTAGGGTTAGGGTTAGGG; Exiqon). After each hybridization, slides were washed three times in 2x SSC at room temperature for 10 min. Both for FISH and CO-FISH, DNA was counterstained with 100 ng/ml DAPI (Sigma-Aldrich) in 1x PBS and slides were mounted in Vectashield (Vectorlabs). Images were acquired with a Zeiss Cell Observer equipped with a cooled Axiocam 506?m camera and a 63X/1.4NA oil DIC M27 PlanApo N objective. Image analysis was performed using ImageJ and Photoshop software.

### EdU incorporation and detection at telomeres

Cells grown on coverslips were incubated in medium containing 2 mM Thymidine (Sigma-Aldrich) for 21 h before replacement with fresh dox-containing medium. After 4 h, 10 μM RO-3306 (Selleckchem) was added and 18 h later 10 μM EdU (Thermo Fisher Scientific) was added to the culture medium, followed by a 2.5 h incubation. Cells were hybridized as for DNA FISH, washed twice with 1x PBS and EdU was detected using the Click-iT EdU Alexa Fluor 488 Imaging Kit (Thermo Fisher Scientific) according to the manufacturer’s instructions. DNA was counterstained with 100 ng/ml DAPI in 1x PBS and coverslips were mounted on slides in Vectashield. Image acquisition and analysis were as for DNA FISH.

### Indirect immunofluorescence (IF)

For HA detection, cells grown on coverslips were incubated in 100% Methanol (Merk) at -20°C for 15 min. For all other IF experiments, cells were incubated in CSK buffer (100?mM NaCl, 300?mM sucrose, 3?mM MgCl_2_, 0.5% Triton X-100, 10?mM PIPES pH 7) for 7 min on ice, fixed with 4% formaldehyde (Sigma-Aldrich) in 1x PBS for 10 min and permeabilized again with CSK buffer for 5 min. Fixed cells were incubated in blocking solution (0.5% BSA, 0.1% Tween-20 in 1x PBS) for 1 h followed by incubation in blocking solution containing primary antibodies for 1 h, three washes with 0.1% Tween-20 in 1x PBS for 10 min each, and incubation with secondary antibodies diluted in blocking solution for 50 min. For combined IF and DNA FISH, cells were again fixed with 4% formaldehyde in 1x PBS for 10 min, washed three times with 1x PBS, incubated in 10 mM Tris-HCl pH 7.2 for 5 min and then denatured and hybridized with a PNA probe (5’-AF568-OO-CCCTAACCCTAACCCTAA-3’; Panagene) as for DNA FISH. DNA was counterstained with 100 ng/ml DAPI in 1x PBS or in 0.1 M Tris-HCl pH 7.2, 0.15 M NaCl, 0.08% Tween-20. Coverslips were mounted on slides in Vectashield. The following primary antibodies were used: a rabbit monoclonal anti-HA (Cell Signaling, 3724; 1:1000 dilution), a rabbit polyclonal anti-RAP1 (Bethyl, A300-306A; 1:2000), a mouse monoclonal anti-TRF2 (Millipore, 05-521; 1:2000), a mouse monoclonal anti-POLD3 (Novus Biologicals, H00010714-M01; 1:500), a mouse monoclonal anti-PML (Santa Cruz Biotechnology, sc-966; 1:500), a rabbit polyclonal anti-pSer33 (Bethyl, A300-246A; 1:2000), a rabbit polyclonal anti-RPA32 (Bethyl, A300-244A; 1:1000), a mouse monoclonal anti-γH2AX (Millipore, 05-636; 1:2000). Secondary antibodies were Alexa Fluor 568-conjugated donkey anti-rabbit IgGs (Thermo Fisher Scientific, A10042; 1:1000) and Alexa Fluor 488-conjugated donkey anti-mouse IgGs (Thermo Fisher Scientific, A21202; 1.1000). Image acquisition and analysis were as for DNA FISH.

### Chromatin immunoprecipitation (ChIP)

Cells were harvested by trypsinization, centrifuged at 500 g at 4°C for 5 min and resuspended in 1% formaldehyde (Sigma-Aldrich) for 20 min at room temperature, followed by quenching with 125 mM glycine (VWR) for 5 min. Cross-linked cells were centrifuged as above and pellets were resuspended in ChIP lysis buffer (1% SDS, 10 mM EDTA, 50 mM Tris-HCl pH 8), sonicated using a Bioruptor apparatus (Diagenode) and centrifuged at 16000 g for 10 minutes at 4°C. 1 mg of lysate was diluted in ChIP dilution buffer (150 mM NaCl, 20 mM Tris-HCl pH 8, 1% Triton X-100, 2 mM EDTA) and incubated with 2 μg of a rabbit monoclonal anti-HA antibody (Cell Signaling, 3724) for 2 h at room temperature. Immunocomplexes were isolated by incubation with Protein A/G PLUS-Agarose beads (Santa Cruz Biotechnology) at 4°C overnight on a rotating wheel. Beads were washed 4 times in ChIP wash buffer (150 mM NaCl, 20 mM Tris-HCl pH 8, 1% Triton X-100, 0.1% SDS, 2 mM EDTA) and once in ChIP final wash buffer (500 mM NaCl, 20 mM Tris-HCl pH 8, 1% Triton X-100, 0.1% SDS, 2 mM EDTA). Beads were incubated in ChIP elution buffer (1% SDS, 100 mM NaHCO_3_) containing 40 μg/ml RNase A (Sigma-Aldrich) for 1 hour at 37°C and DNA was extracted using the Wizard SV gel and PCR cleanup system (Promega). Input and immunoprecipitated DNA was subjected to quantitative PCR using the oligonucleotides shown in Table 1. QPCRs were performed using the iTaq Universal SYBR Green Supermix (Bio-Rad) on a Rotor-Gene Q (Qiagen) instrument with a 2-step program (45 cycles of denaturation at 95°C for 15 sec, annealing and extension at 60°C for 30 sec). Data analysis was performed using the Rotor-Gene 6000 Series Software 1.7.

**Table 1:**
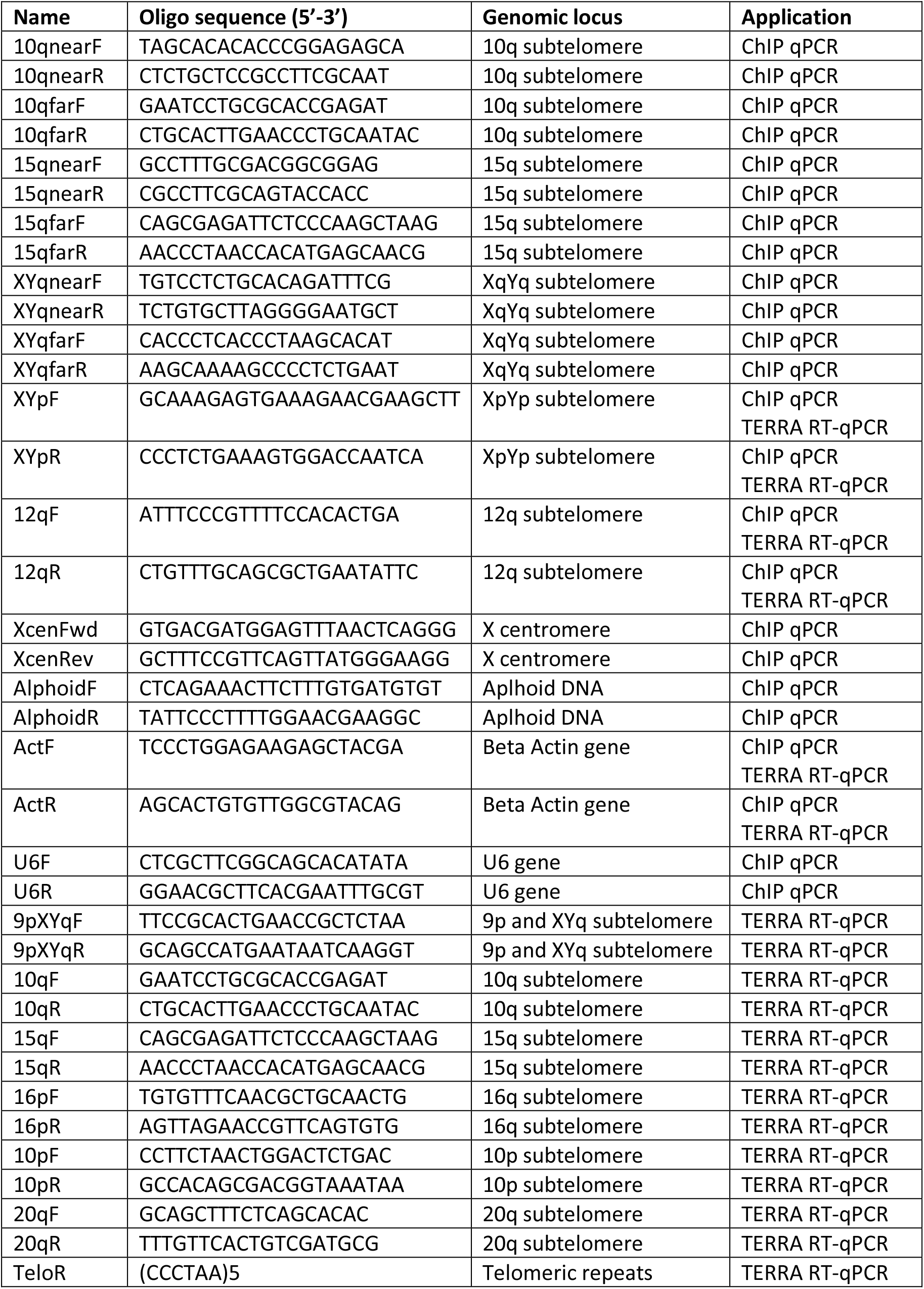
oligonucleotides used in this study.

### Reverse transcription and quantitative PCR

Total RNA was isolated using the TRIzol reagent (Thermo Fisher Scientific) followed by chloroform extraction and treated three times with 3.5 U of DNaseI (Qiagen) for 45 minutes at room temperature. 5 μg of RNA were reverse transcribed with 0.5 μM TeloR and 0.05 μM ActinR oligonucleotides (Table 1) and Superscript III (Thermo Fisher Scientific) according to the manufacturer’s instructions. Quantitative PCRs were performed and analyzed as for ChIP using the oligonucleotides shown in Table 1. Actin values were used as normalizers.

### C-circle assay

Genomic DNA was isolated by phenol:chloroform extraction and treatment with 40 μg/ml RNaseA (Sigma-Aldrich), followed by ethanol precipitation. Reconstituted DNA was digested with HinfI and RsaI (New England Biolabs) and again purified by phenol:chloroform extraction. 500 ng of digested DNA were incubated with 7.5 U of phi29 DNA polymerase (New England Biolabs) in presence of dATP, dTTP and dGTP (1 mM each) at 30°C for 8 h, followed by heat-inactivation at 65°C for 20 min. Amplification products were dot-blotted onto nylon membranes (GE Healthcare) and hybridized at 55°C overnight with a double-stranded telomeric probe (Telo2 probe), radioactively labeled using Klenow fragment (New England Biolabs) and [α-32P]dCTP. Post-hybridization washes were twice in 2x SSC, 0.2% SDS for 20 min and once in 0.2x SSC, 0.2% SDS for 30 min at 50°C. Radioactive signals were detected using a Typhoon FLA 9000 imager (GE Healthcare) and quantified using ImageJ software.

### Statistical analysis

For direct comparison of two groups, we employed a paired two-tailed student’s t-test using Microsoft Excel or a nonparametric two-tailed Mann-Whitney U test using GraphPad Prism. Values are indicated as: \**P*?<?0.05, \*\**P*?<?0.005, \*\*\**P*?<?0.001, \*\*\*\**P*?<?0.0001.

## Supplementary information

### SUPPLEMENTARY FIGURES

**Figure S1:**
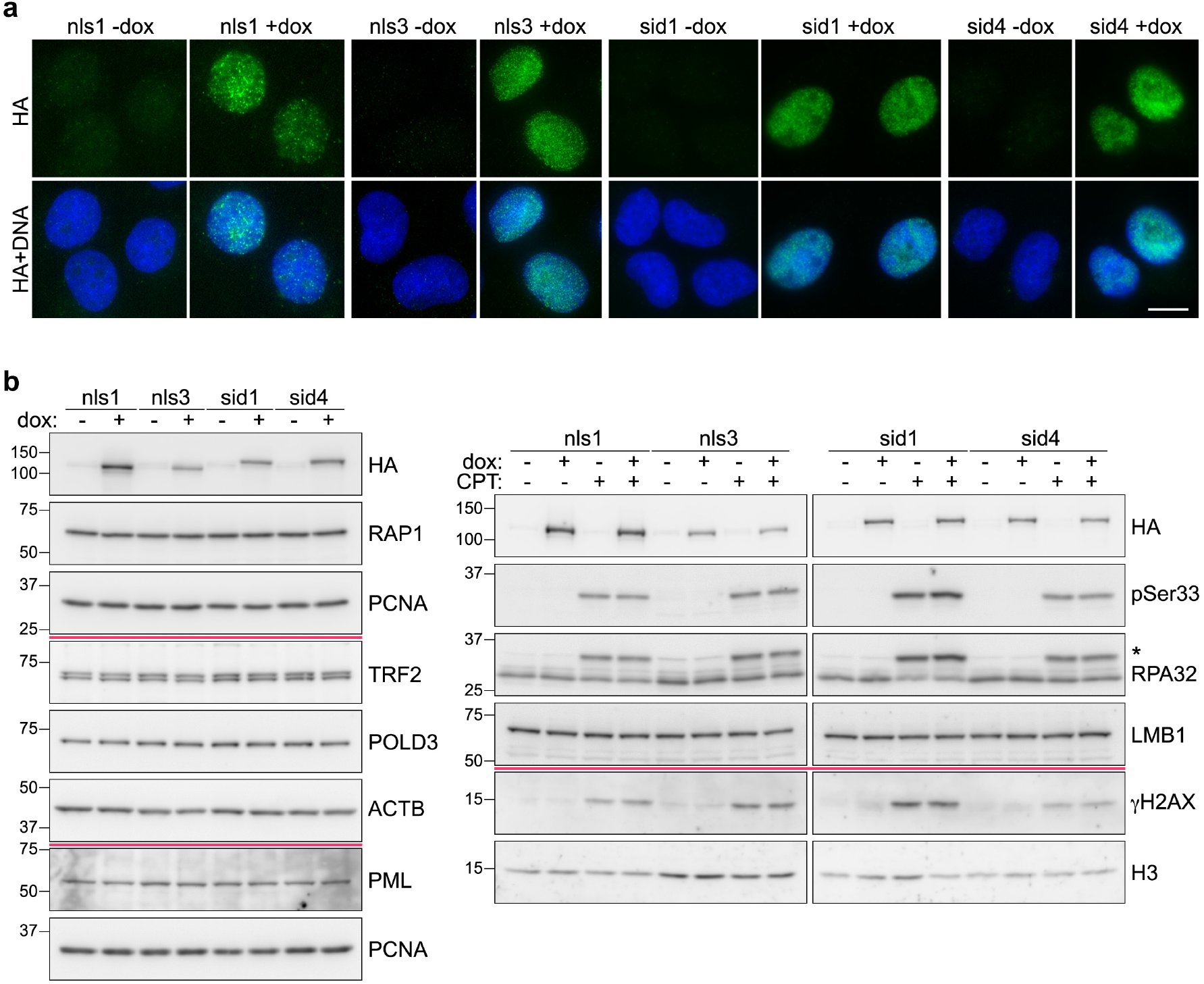
Expression of T-TALEs and endogenous proteins used in this study. (**a**) Examples of anti-HA tag IF (green) in the indicated cell lines treated with dox for 24 hours or left untreated. DAPI stained DNA is in blue. Scale bar: 10 μm. (**b**) Western blot analysis of the indicated cell lines treated with dox as in **a**. Red lines separate signals from different membranes. Beta actin (ACTB), PCNA, Lamin B1 (LMB1) and histone H3 serve as loading control. Numbers on the left are molecular weights in in kDa. CPT: camptothecin.

**Figure S2:**
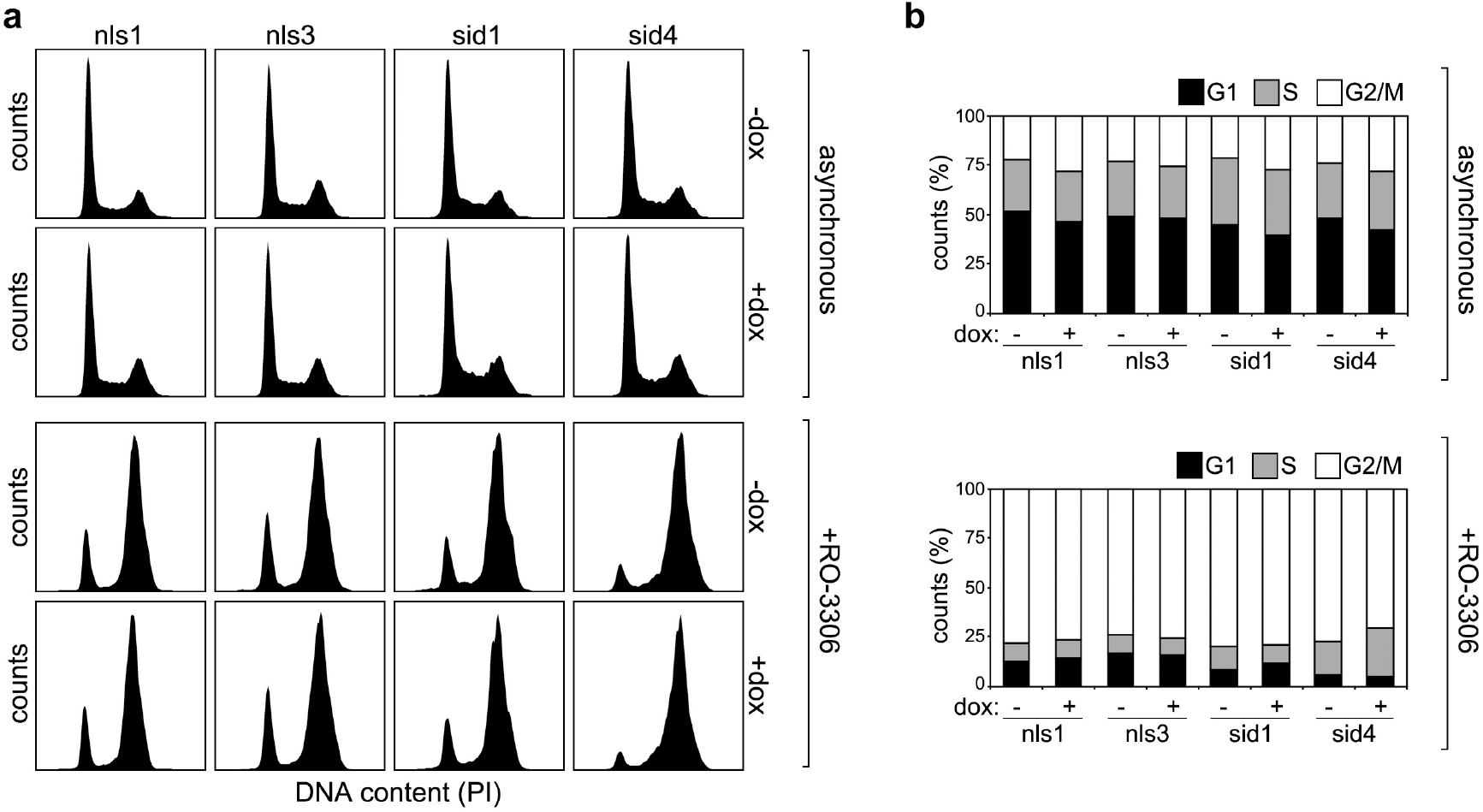
Cell cycle distribution analysis of T-TALE cells. (**b**) FACS profiles of the indicated propidium iodide (PI)-stained cells treated or not with dox and with the CDK1 inhibitor RO-3306. Cell counts (y axis) are plotted against PI intensity (x axis). Cells were harvested after 24 (asynchronous) or 24.5 (RO-3306-treated) hours of dox treatment (see methods for details). (**b**) Quantifications of experiments as in **a**. The graphs show the percentage of cells in G1, S and G2/M phases from one representative experiment.

**Figure S3:**
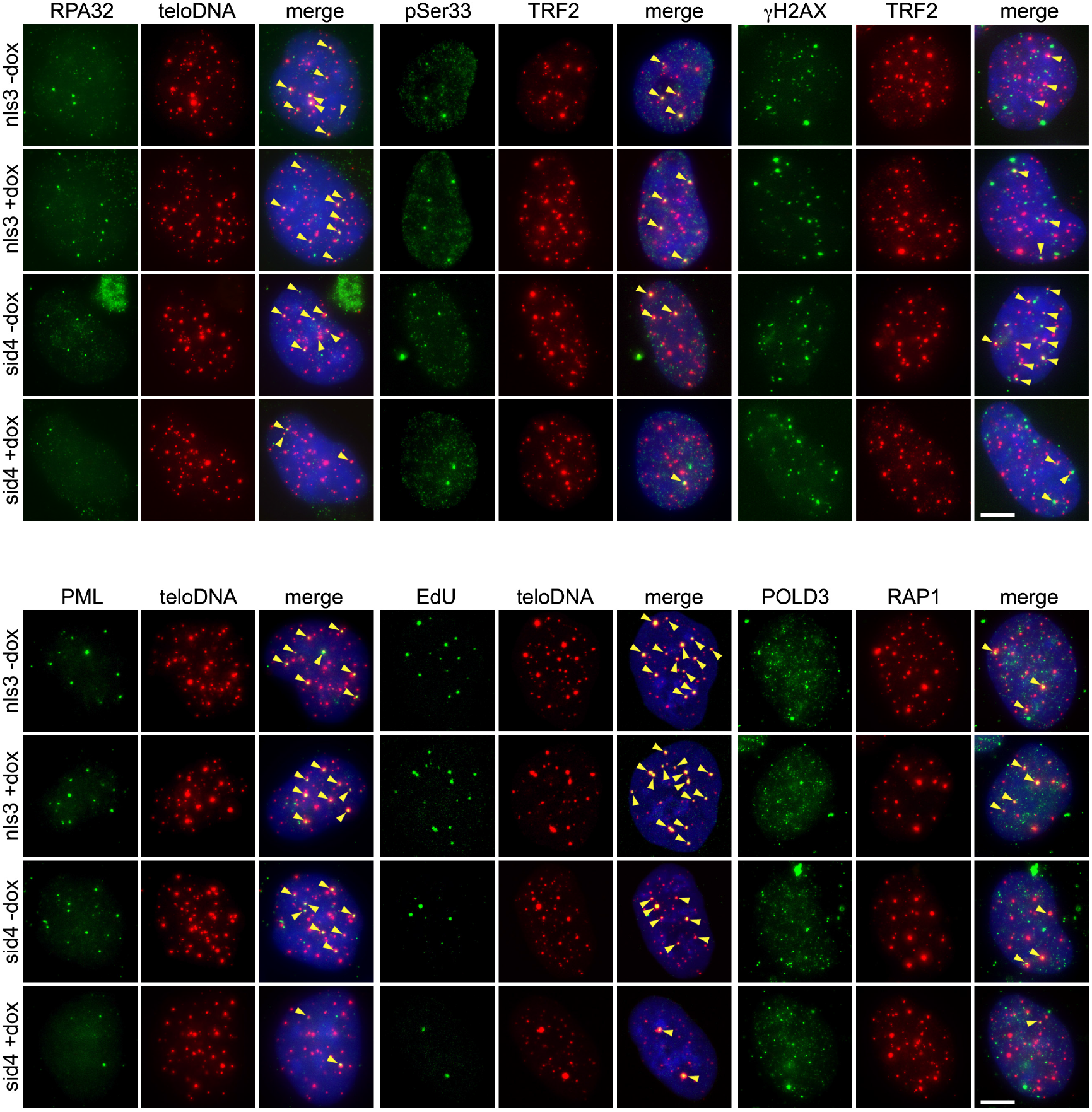
TERRA transcription inhibition alleviates telomere instability and ALT activity. Examples of experiments as in Figures 2 and 3 performed in nls3 and sid4 cells. Markers and DAPI stained DNA are shown with the same colors as in Figures 2 and 3. Arrowheads in the merge panels point to co-localization events. Scale bars: 5 μm.

**Figure S4:**
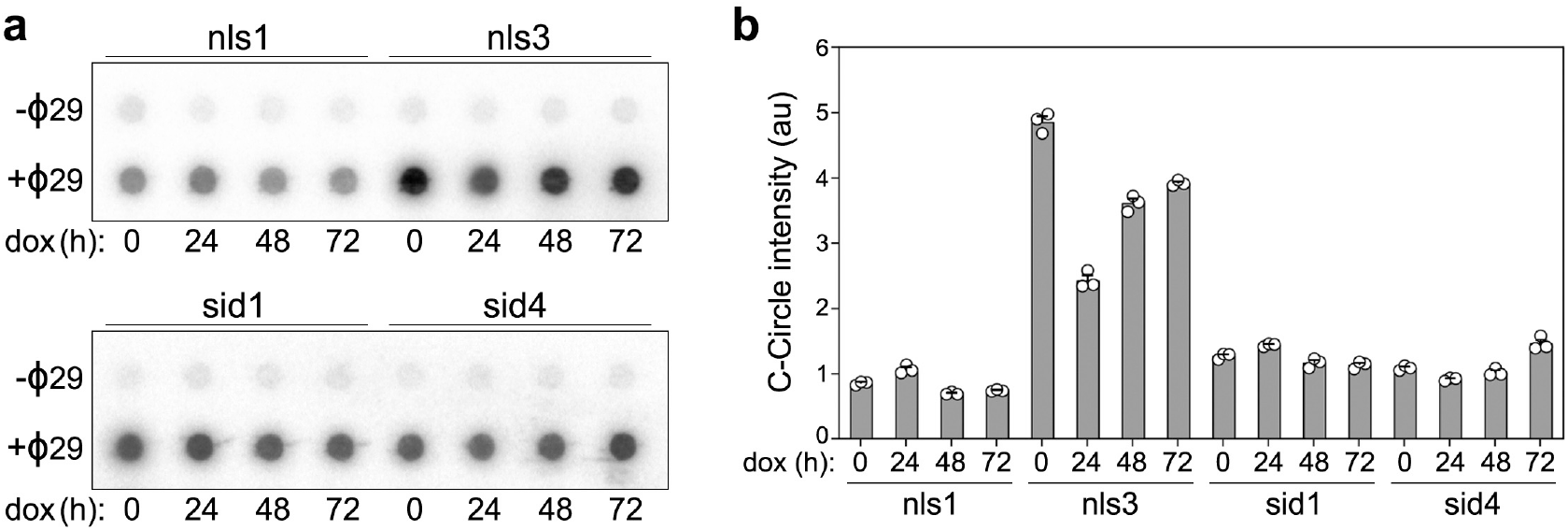
C-circle analysis in T-TALE cells. (**a**) C-circle assay analysis of genomic DNA from the indicated cells treated with dox for up to 72 hours. Reaction products were dot-blotted and hybridized to a radiolabeled telomeric probe. Control reactions were performed in absence of phi29 polymerase (-Φ29). (**b**) Quantifications of C-circle signals from experiments as in **a**. Bars and error bars are means and SEMs from 3 independent experiments. Circles are single data points.

